# Search for and function of different genes in phloem and xylem of poplar

**DOI:** 10.1101/2020.01.20.912493

**Authors:** Xiaoyan Li, Xiangning Jiang

## Abstract

We used the microarray data of poplar gene in the laboratory and the GPDNN model to preprocess the data and conduct the median adjustment, and then studied and analyzed the phloem and xylem genes with good clustering effect. Among them, a gene that was significantly different and related to plant hormones was selected for analysis. Firstly, the function of this gene was predicted to be auxin response factor. All genes of auxin response factor family were selected from poplar transcription factor database for evolutionary and homologous affinity analysis, and the relationship with homology was analyzed according to the expression semaphore correlation between genes. Finally, the number of up-regulated, down-regulated and fine-tuned genes in the tissues of budding, root, xylem and phloem was selected by using the median, and the relationship between the number of up-regulated genes and their semaphores and auxin content in the tissues was analyzed.

## 1 Introduction

Life phenomenon is one of the miracles of nature, heredity is an important content in life. Since 1866, when the father of genetics (Mendel) proposed genetic factors, human beings have realized the importance of heredity. The human genome project (HGP) made human beings understand themselves. With the completion of HGP sequencing, scientists were faced with tens of thousands of genes, and the functions of thousands of genes needed to be studied simultaneously, and the complex network of gene expression and regulation also needed to be studied in parallel. Gene chip technology is characterized by high throughput [1], making it widely used in many fields such as gene expression analysis, disease diagnosis and treatment.

In this paper, based on the poplar genome chip obtained on the basis of molecular biology, GPDNN[2] data processing method was used to preprocess the poplar genome chip data in our laboratory. After cluster analysis and evaluation [3], the phloem and xylem genes with good clustering effect were finally selected for further analysis. First, Wilcox [4]was used to detect phloem and xylem genes, and the differentially expressed genes were selected when p-value<0.05. We selected a gene with significant differences, which is related to auxin according to the GO process marked in Affymetrix marker document [1]. Through our prediction of gene function, this gene is auxin response factor. We found 37 gene models of ARF gene in poplar transcription factor database, selected 32 gene models represented by probes, and conducted homologous evolutionary analysis on them. Through gene expression correlation, we built a network to analyze the relationship between gene expression correlation and homology. We used the corresponding gene semaphore to select the up-regulated, down-regulated and fine-tuned gene distribution in bud, root, xylem and phloem according to the median. Finally, the relationship between the number of up-regulated genes and the number of signals and the content of auxin in different tissues was analyzed according to the content of auxin in different tissues detected by internal standard ^13^C_6_-IAA method.

### 1.1 Biological molecular basis

From the late 1950s to the 1960s, the “central law” and the theory of operon were successively proposed, which successfully cracked the genetic code and clarified the mechanism of the flow and expression of genetic information[5]. DNA and RNA are important substances in biological inheritance. They are composed of four kinds of deoxyribonucleic acid, which are Adenine, Guanine, Thymine and Cytosine. RNA has no thymine T and contains Uracil. Purines pair with pyrimidines in DNA, A pairs with T, C pairs with G, and A pairs with U in RNA. The sequence of nucleotides in DNA molecules not only determines the basic structure of all RNA and proteins in the cell, but also indirectly controls the production, operation and function of all active components in the cell through the function of proteins (enzymes).

The central law of eukaryotic biology [5] mainly describes the important law of interaction between DNA, RNA and protein, which mainly includes three processes: DNA replication; DNA is transcribed to form RNA; RNA is translated into proteins. DNA replicates itself in the nucleus, and then is transcribed to form nuclear non-uniform RNA. After cutting, splicing and other processes, mature mRNA is formed. Subsequent studies have found that RNA can be reversely transcribed into DNA to guide protein synthesis, a finding that complements the central principle. With the development of biological individuals, DNA molecules can sequentially transform the genetic information they carry into proteins through the codon - anticodon subsystem, performing various physiological and biochemical functions.

### 1.2 Gene chip

In the early 1990s, Affymetrix, an American company, produced the world’s first oligonucleotide chip [6] by using in situ synthesis of oligonucleotides. Thousands of densely arranged molecular microarrays integrated on the chip can analyze a large number of biomolecules in a short period of time, enabling people to quickly and accurately obtain the biological information in samples, which is hundreds and thousands of times more efficient than traditional detection methods and convenient for modern researches on a large number of genes [7].

The design principle of oligonucleotide chip is introduced as follows:

Oligonucleotide chip is based on the principle of reverse hybridization, the design and synthesis of good dozen beforehand to dozens of bases of oligonucleotides through sample points at a fixed onto the glass, or by in situ synthesis technology of fixed on the glass, and fluorescent tags for sequencing column under certain conditions, after washing the scan for monitoring information. Oligonucleotide chips usually use in situ synthesis, which combines solid-phase DNA synthesis with photolithography. Technical principles [1] 1: computer chip substrate;2: activate the computer chip surface; 3: light inactivation “A” mask;4: cross-linked with adenosine (A) reagent; 5. Repeated synthesis cycle, in which a photosensitive protective group is attached to the end of the 5 ‘-hydroxyl group of the synthetic base monomer.

Gene chip experiments generally include the following steps [1]:

1. Chip probe design. According to the needs of experimental purposes, the probe complementary to the target sequence was synthesized and fixed to the carrier by means of the hybrid complementary principle. The sensitivity and specificity of gene chip hybridization is the core of chip technology, and Affymetrix has developed a unique pm-mm probe scheme. Each probe set for each gene on the chip consists of 10-20 probe pairs, each consisting of two probe units, one of which is a perfectmatch (PM) and the other a mismatch (MM) of the 13th base in the middle of the sequence.
2. Sample preparation. Total RNA or purified poly-a mRNA was extracted from tissues or cells to synthesize double-stranded cDNA, and then in vitro transcription was used to synthesize cRNA, which was fragmented before hybridization.
3. Hybridization of sample and probe, cleaning and coloring of probe. The obtained samples were incubated with the prepared probe for 16 hours to ensure the complete hybridization. After hybridization, the samples were combined with complementary probes, and the uncombined samples were rinsed out before scanning to ensure the accuracy of scanning data.
4. Scan to obtain data. The laser scanner was used to scan the hybrid microarray and obtain the fluorescence intensity of the labeled hybrid sequence. The fluorescence intensity obtained was the required data.
5. Data analysis. The obtained data can be preprocessed in a certain way so that the obtained data can be trusted for future use.

### 1.3 Data processing

After the chip is scanned by laser, the output is calculated according to the fluorescence intensity and becomes the chip data. However, the chip data has many noise points, such as background and other factors, which affect the accuracy of data evaluation. For this reason, in the following period of time, many bioinformatists have done a lot of calibration work on biochip data. Cheng Li[9]’s statistical model, Irizarry’s RMA model [10]and Li Zhang’s PDNN model [11]are all mathematical models for the preprocessing of chip data. Ning Jiang[12] et al., in BMC Bioinformatics in 2008, made a lot of comments on the above correction pretreatment methods, and analyzed that PDNN was a better one of the above data pretreatment methods. Unfortunately, the Mismatch (MM) data was not taken into account in the PDNN data processing model, which was known to play an important role in gene expression level based on the chip principle. The GPDNN model established by wei wei of Peking University [2]makes up for this defect and makes data processing more reliable. We will use GPDNN model to preprocess poplar chip data and evaluate the data according to the clustering effect.

PDNN model divides hybridization into specific hybridization and non-specific hybridization. Specific hybridization refers to the binding of the probe to the target sequence, while non-specific hybridization refers to the binding of the probe to other sequences. Many studies have found that the size of the intensity of hybridization by combining ability (binding affinity), two relations can be absorbed by the Langmuir theory (Langmuir Adsorption Model). Among the 25 base-long probes, different positions of the probes have different effects on the binding ability. The edge part has less attraction and fixation ability to the target sequence than the middle part. The binding ability is affected by free energy. According to the study of Wang[13] [], for probe sequences with high T and C content, its fluorescence intensity is higher, which indicates that the probe sequences themselves also affect the ability of binding target sequences. Li Zhang[11] proposed in the model that the binding energy was determined by two factors: position and stacking free energy of base sequence, and assumed that the stacking energy was determined by the adjacent base pair. The combining capacity is the linear sum of the position weights and the stacking energy. However, because the non-specificity contains certain information, the PDNN model ignores this point.

Wei wei improved the original PDNN algorithm with the following specific methods: first, Wilcoxon[4] [] symbol test was conducted to determine two training data sets. Then the MM probe information was added to the non-specific parameter estimation model, and the non-specific binding and specific binding parameters were estimated respectively.[see appendix for specific GPDNN algorithm model]

## Materials and methods

### 2.1 Experimental materials

Our poplar chip is an Expression Microarray made by Affymetrix. Poplar expression profile chip is specially designed to control gene expression. Poplar gene expression profile chip contains more than 61,000 probe groups, representing more than 56,000 transcripts and gene products [1] [].Poplar chip design was conducted by Affymetrix chip design team in cooperation with poplar research authority. The genes in the chip design process were selected from UniGene builds, mRNAs and ESTs in GenBank of all poplar species, ensuring the integrity of the design [1] [].

Our poplar microarray consists of three biological replicates at the time level (spring, summer, autumn, winter) and tissue level (root, stem, xylem cambium, phloem cambium, leaf), with a total of 60 microchips. But for some special reason, four chips were not made successfully. So now we’re left with 56 chips. The following is described in mathematical language:

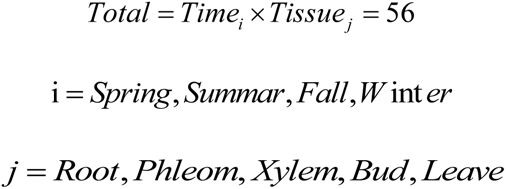

### 2.2 methods screening of differential genes

We evaluated the clustering effect of the preprocessed data, and the results showed that the gene clustering effect between phloem and xylem was better [figure]. For this reason, phloem and xylem were selected for future research and analysis.

First, we need to pick out the genes that differ between the phloem and xylem. At present, there have been many studies on the methods of selecting differential genes, such as Subramanian’s GSEA (Gene sets enrich analysis) [14], etc., and the screening methods of these differential genes are based on a large number of data.Our poplar chip phloem and xylem data are less, so the main experimental method to find the different genes is wilcoxon non-parametric test [4]. The non-parametric test is a test that is independent of the population distribution. It does not depend on the form of the population distribution. The non-parametric test is essentially a test to see if the position (median) of the population distribution is the same.Wilcoxon nonparametric test method is an improved symbol test. By arranging observed values in order from small to large, rank order is made, rank is obtained and hypothesis is tested. We first made the statistical hypothesis that the median expression in phloem was equal to the median expression in xylem, and the alternative hypothesis was in the opposite direction, that the median expression in phloem was not equal to the median expression in xylem. Then R software [15] was used to make relevant calculations of wilcox.test, and then the genes with significant differences (that is, P <0.05) were screened out according to p-value, and finally the corresponding annotation of GO[16] was found. Our statistical assumptions are:

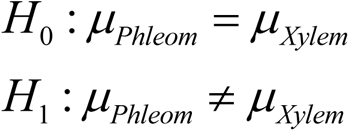

When p-value <0.05 was selected, the genes with significant difference were the differentially expressed genes in phloem and xylem.

## 3 results and discussion

### 3.1 data preprocessing and clustering analysis

After the poplar chip data were preprocessed by the GPDNN model and then the median leveling was carried out, we conducted cluster analysis and evaluation on the 45 chip data [3], which did not include the leaf data. From figure 3.1, it can be concluded that the phloem and xylem genes of poplar can cluster into different species according to the seasons of spring, summer, autumn and winter respectively, that is, the clustering effect of the phloem and xylem genes of poplar is good. Therefore, poplar phloem and xylem genes were selected for the following analysis.

**Figure 3.1.**
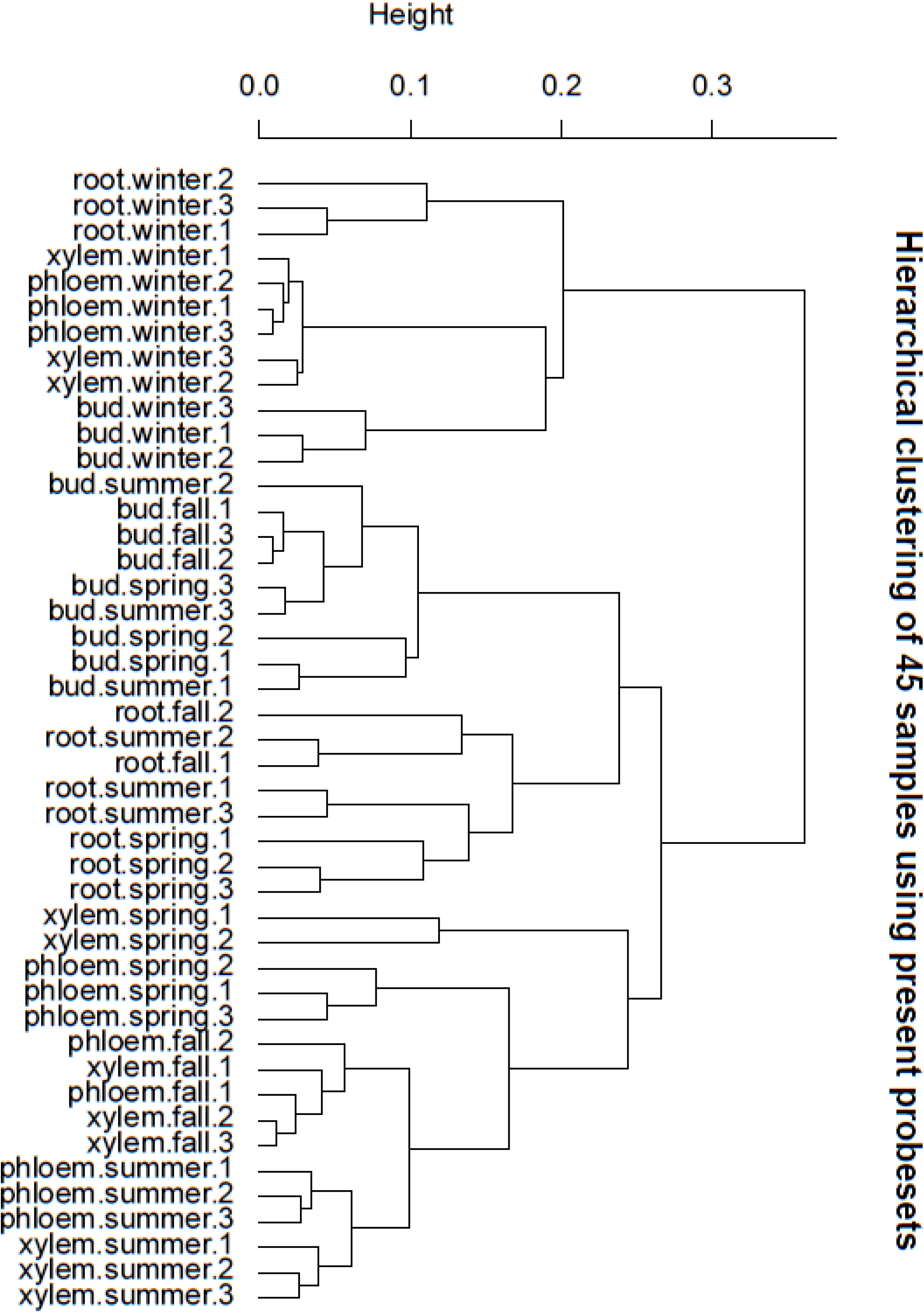
the result of cluster analysis

### 3.2 selection of phloem and xylem genes

For phloem and xylem genes, we used R software [15] to calculate the wilcoxo. Test and extracted the final p-value. After sorting by p-value, we found the corresponding annotation of the respective GO[16] process. We selected the genes with significant differences between phloem and xylem when p<0.05 after calculation. Finally, we found a total of 3,239 genes with significant differences, of which 1,039 had corresponding GO annotation. These genes are classified according to the processes they are involved in, as shown in figure []. Among them, 103 genes are involved in protein phosphorylation, 83 genes are involved in metabolism, 69 genes are involved in DNA transcription regulation, 67 genes are involved in transcription regulation and so on. It can be concluded from the results that most of the differential genes are involved in the basic metabolic process of poplar.

**figure 3.2:**
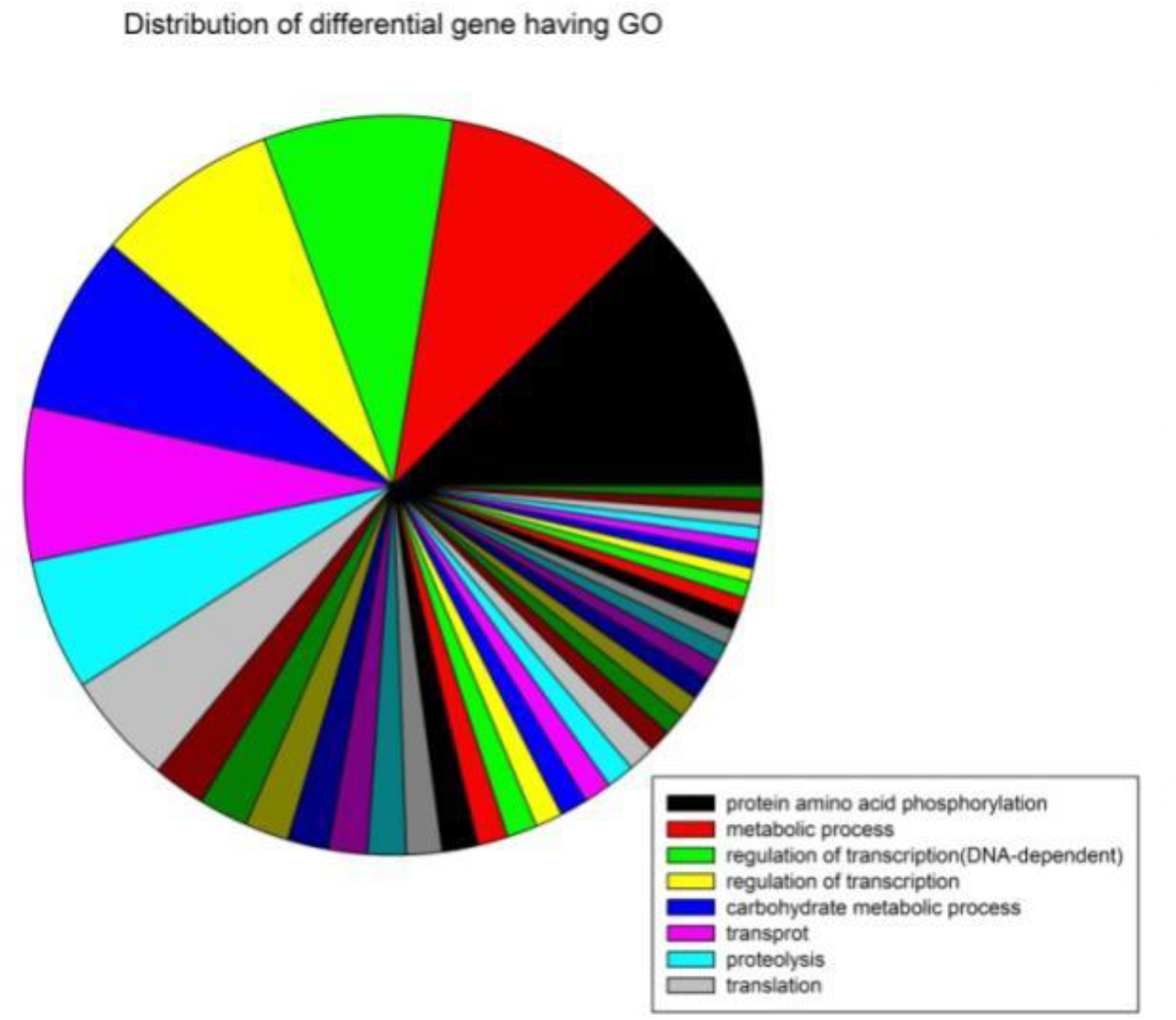
Distribution of differential gene having GO

**figure 3.3.**
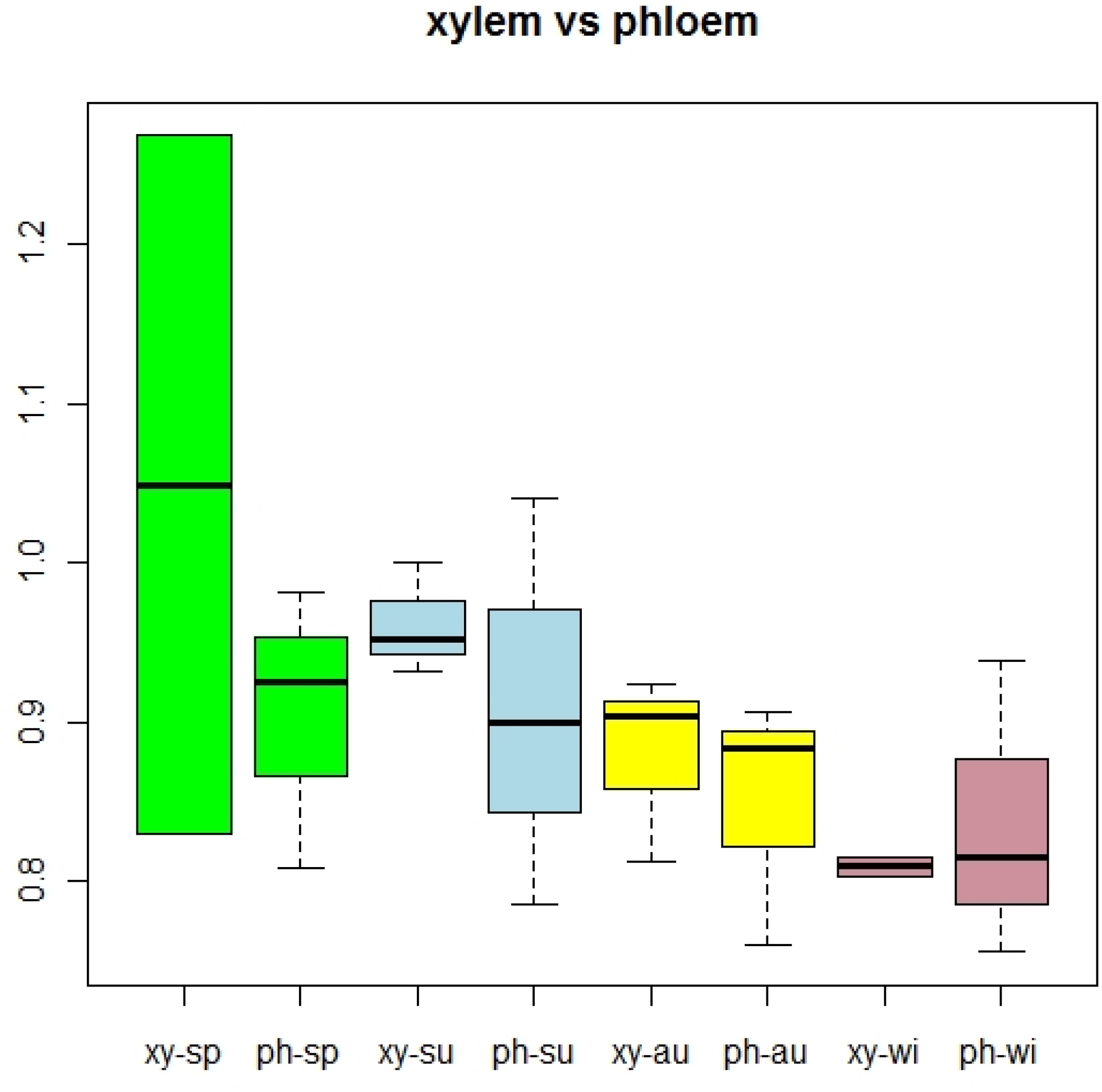
shows the differential expressions of phloem and xylem in spring, summer, autumn and winter

In addition, we also used R for the differential expression of poplar phloem and xylem to make box diagram:

1. According to the seasonal standard, the xylem gene expression in spring, summer and autumn was higher than that in phloem, while the phloem gene expression in winter was not significantly different from that in xylem. In spring, summer and autumn, xylem gene expression was significantly higher than phloem in spring, followed by summer, and the difference between xylem gene expression and phloem gene expression in autumn was the smallest.
2. Tissue as the standard: in xylem and phloem, with the changes of spring, summer, autumn, winter and four seasons, gene expression levels in each tissue showed a declining trend. In winter, the gene expression level was almost the same in all tissues, while in winter, the gene expression level in all tissues was very low when the trees were dormant and overwintering.

We list 10 genes with large p_value(Table 3.1):

**Table 3.1.**
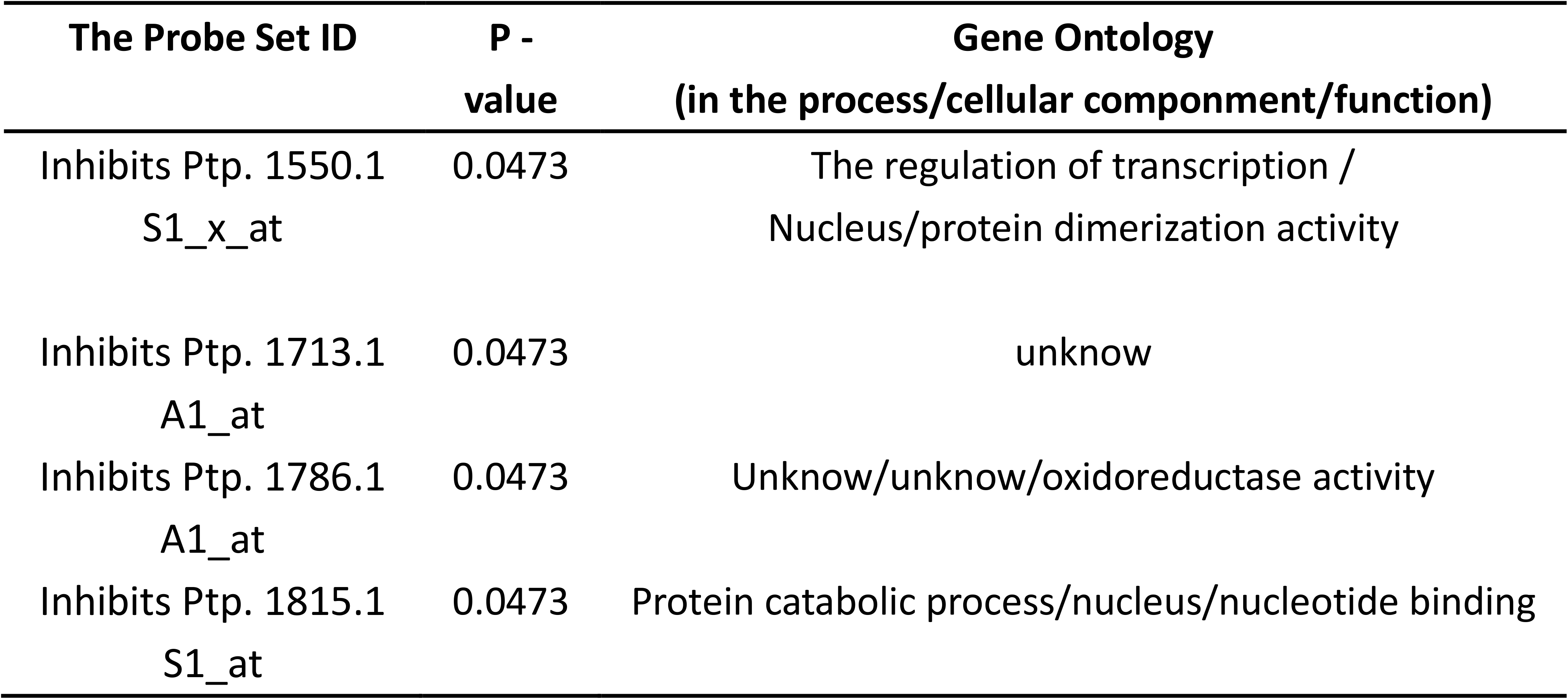
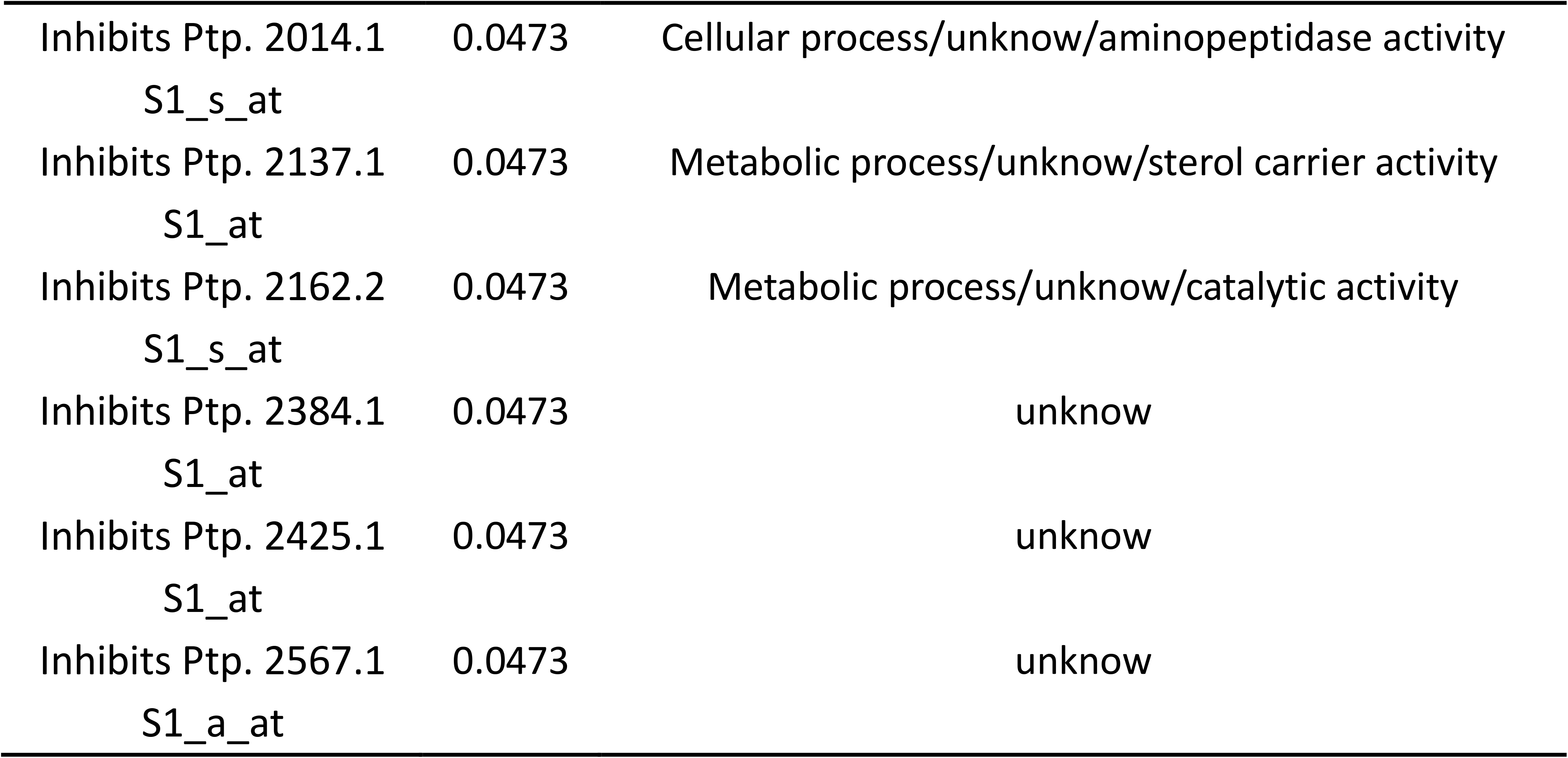
list 10 genes with large p_value

From the genes with significant differential expression (i.e., p-value <0.05), Wilcox [4] was first used for gene expression detection, and then an auxin related gene was selected as the target gene for research and analysis. According to the process of GO[16], the function of this purpose gene(Table 3.2) is to respond to plant hormones, and the encoded protein is located in the nucleus, mainly through binding to DNA.

**Table 3.2.** The function of this purpose gene

| The Probe Set ID | P - value | Gene Ontology<br>(in the process/cellular component/function) |
| --- | --- | --- |
| PtpAffx. 205505.1 S1_at | 0.039975 | Response to hormone stimulus/nucleus/DNA binding |

### 3.3 function prediction of target genes

A total of 1049bp of matching genetic information was found from the data provided by Affymatrix (appendix). First, complete nucleic acid comparison was conducted between the gene sequence of the target gene and the poplar genome data in NCBI database [19]. The results are shown in figure 3.4.

**Figure 3.4.**
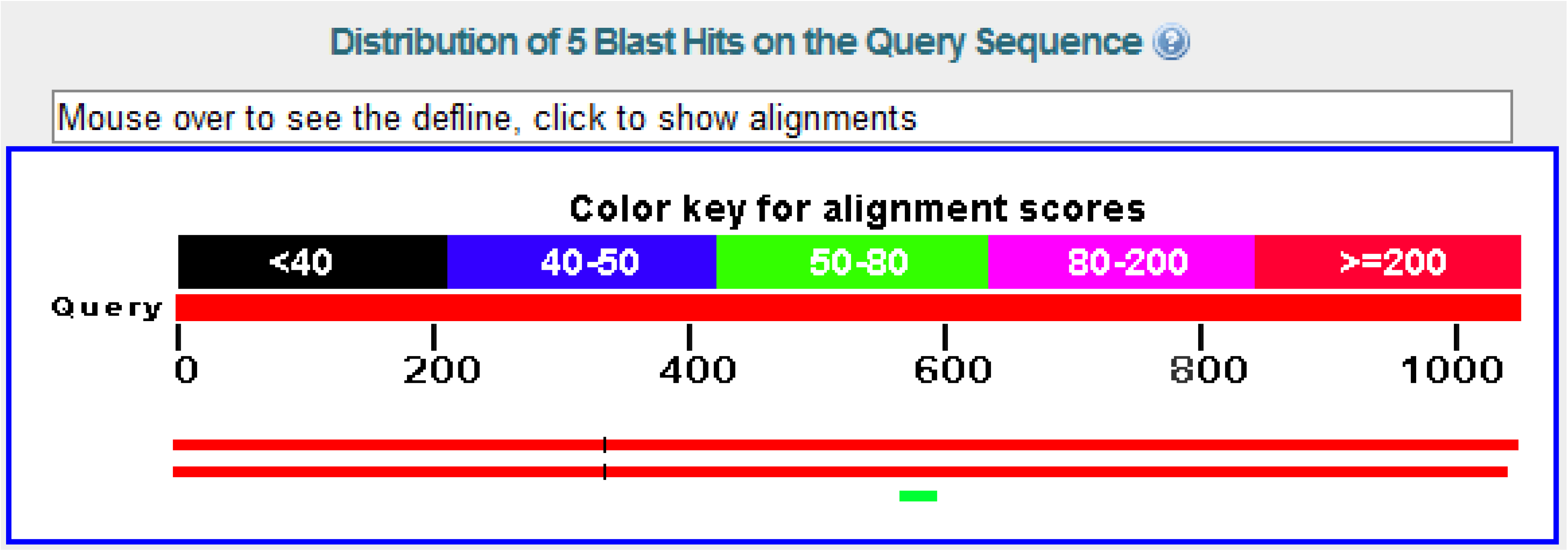
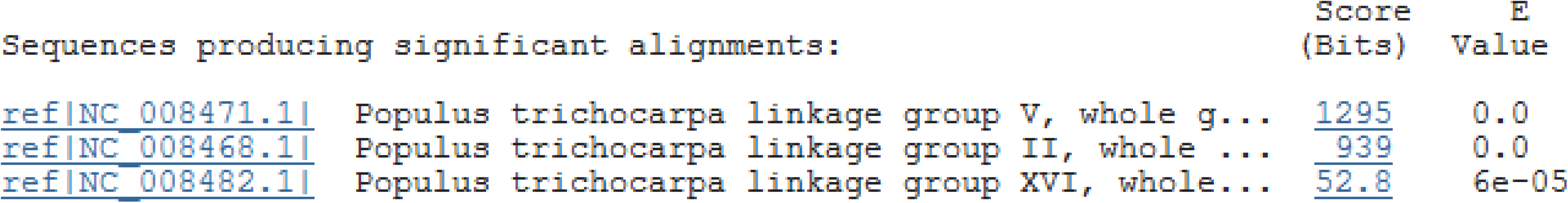
The result of Blast

We choose high grade poplar chain group to predict, will first match gene location and extension 1000 bp respectively, extended full genetic 4717 bp (appendix), and online prediction of the gene prediction software softberry, and finally the complete gene prediction, as shown in figure3.5. As shown then we estimated by softberry protein sequence in the Pfam database, to protein function prediction results show that both in the Pfam-A more accurate results(Table 3.3). From the results, we selected the gene function with high score and predicted the gene function as Auxin response factor (ARF).

**table 3.3:** functions of pfam predictive genes

| Family | The Description | The HMM |  | Bit Score | E - the value |
| --- | --- | --- | --- | --- | --- |
|  |  | The From | The To |  |  |
| B3 | B3 DNA binding domain | 1 | 95 | 82.8 | 8.6 e-24 |
| Auxin_resp | Auxin response factor | 1 | 83 | 85.1 | 1.8 e-24 |

**figure 3.5:**
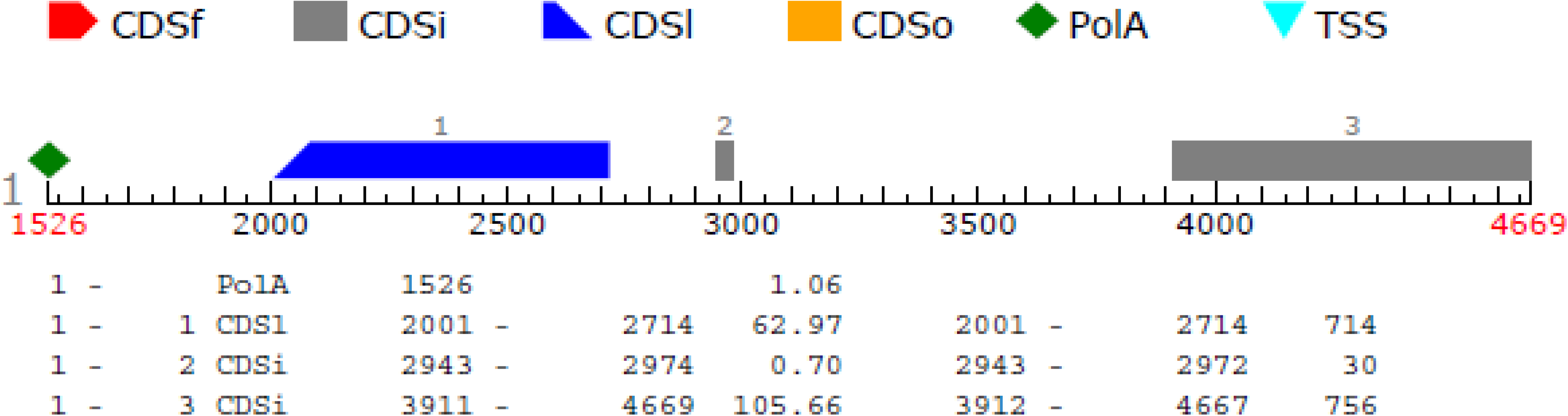
prediction of gene integrity after expansion

### 3.4 analysis of aspen auxin response factor family (ARFs)

#### 3.4.1 ARFs

The ARF family is a class of nuclear proteins that act as transcription factors. Similar to Aux/IAA, ARFs also contains four conservative domain structure, respectively called N terminal dna-binding domain (DBD) structure, the middle region (MR), structural domain III and IV (both collectively known as polymerization structure domain (CTD) C terminal 2 [20]. Different domains have different functions, DBD can directly bind AuxREs of auxin regulatory gene promoter, and DBD cannot determine its regulatory characteristics. Auxin inductance with AuxRE promoter depends on the intermediate and c-terminal regions of ARFs. Structure in the end of the ARF and Aux/IAAs C domain III and IV are highly conserved, CTD is mediated Aux/IAAs and ARFs homologous polymerization of polymerization and two different source two area, the combination of the polymerization process to adjust ARFs and AuxREs and auxin response gene expression [21].

We found 37 ARF gene models from the plant transcription factor database [38], and then located the ARF family members according to the auxin response factor model [39] established by Peking University [table 3.4].

**Table 3.4.** 37 ARF gene models from the plant transcription factor database

| NO. | Gene name | Protein ID | Chromosome |
| --- | --- | --- | --- |
| 1 | Fgenes4_pg. C_LG_I002802 | 753199 | LG_I |
| 2 | Gw1. I. 7907.1 | 179307 | LG_I |
| 3 | Gw1. I. 8521.1 | 179921 | LG_I |
| 4 | Gw1. II. 1258.1 | 420624 | LG_II |
| 5 | Gw1. II. 3332.1 | 422698 | LG_II |
| 6 | Gw1. III. 1140.1 | 424738 | LG_III |
| 7 | EstExt_fgenes4_pm. C_LG_III0550 | 830919 | LG_III |
| 8 | Gw1. IV. 3880.1 | 198791 | LG_IV |
| 9 | EstExt_Genewise1_v1. C_LG_IV2935 | 713904 | LG_IV |
| 10 | Fgenes4_pg. C_LG_IX001411 | 768360 | LG_IX |
| 11 | Fgenes4_pg. C_LG_IX001438 | 768387 | LG_IX |
| 12 | EstExt_fgenes4_pg. C_LG_V0901 | 818500 | LG_V |
| 13 | Gw1. V. 808.1 | 205407 | LG_V |
| 14 | Gw1. VI. 2366.1 | 428694 | LG_VI |
| 15 | Fgenes4_pg. C_LG_VIII000301 | 765287 | LG_VIII |
| 16 | EstExt_fgenes4_pm. C_LG_X0888 | 833860 | LG_X |
| 17 | EstExt_Genewise1_v1. C_LG_XI2869 | 727995 | LG_XI |
| 18 | EstExt_fgenes4_pm. C_LG_XII0386 | 834367 | LG_XII |
| 19 | Gw1. XIV. 2727.1 | 245984 | LG_XIV |
| 20 | Gw1. XIV. 424.1 | 243681 | LG_XIV |
| 21 | Gw1. XIV. 1750.1 | 245007 | LG_XIV |
| 22 | eugene3.00150845 | 575450 | LG_XV |
| 23 | EstExt_fgenes4_pm. C_LG_XVI0323 | 835516 | LG_XVI |
| 24 | Fgenes4_pg. C_LG_XVIII000045 | 778677 | LG_XVIII |
| 25 | eugene3.150910001 | 583448 | scaffold_15091 |
| 26 | EstExt_fgenes4_pg. C_1640064 | 828493 | scaffold_164 |
| 27 | Fgenes4_pg. C_scaffold_187000006 | 786899 | scaffold_187 |
| 28 | eugene3.00280060 | 589324 | scaffold_28 |
| 29 | gw1.44.432.1 | 289219 | scaffold_44 |
| 30 | eugene3.00660262 | 595068 | scaffold_66 |
| 31 | eugene3.08470003 | 596587 | scaffold_847 |
| 32 | EstExt_fgenes4_pm. C_860029 | 836829 | scaffold_86 |
| 33 | grail3.0003020302 | 642997 | LG_II |
| 34 | grail3.0002064402 | 652033 | LG_V |
| 35 | EstExt_fgenes4_pg. C_1500013 | 828321 | scaffold_150 |
| 36 | gw1.28.631.1 | 282057 | scaffold_28 |
| 37 | gw1.28.632.1 | 282058 | scaffold_28 |

#### 3.4.2 correlation analysis of ARFs homologous evolution and expression

##### 3.4.2.1 homology and evolutionary analysis of ARFs

In the plant transcription factor database [38], we obtained the corresponding CDS sequence of 37 genes of ARF family, and analyzed and studied the relationship and evolutionary relationship of ARF family by using MEGA4.

From the evolutionary tree [see figure 3.7], we can roughly divide the ARF gene family into three categories (classI, classII, classIII), among which the first category includes two small categories (classIa, classIb), and the homology between classIa and classIb is only 44%.From an evolutionary perspective, the genes in classIII evolved first, followed by classI and classII.

**figure 3.7.**
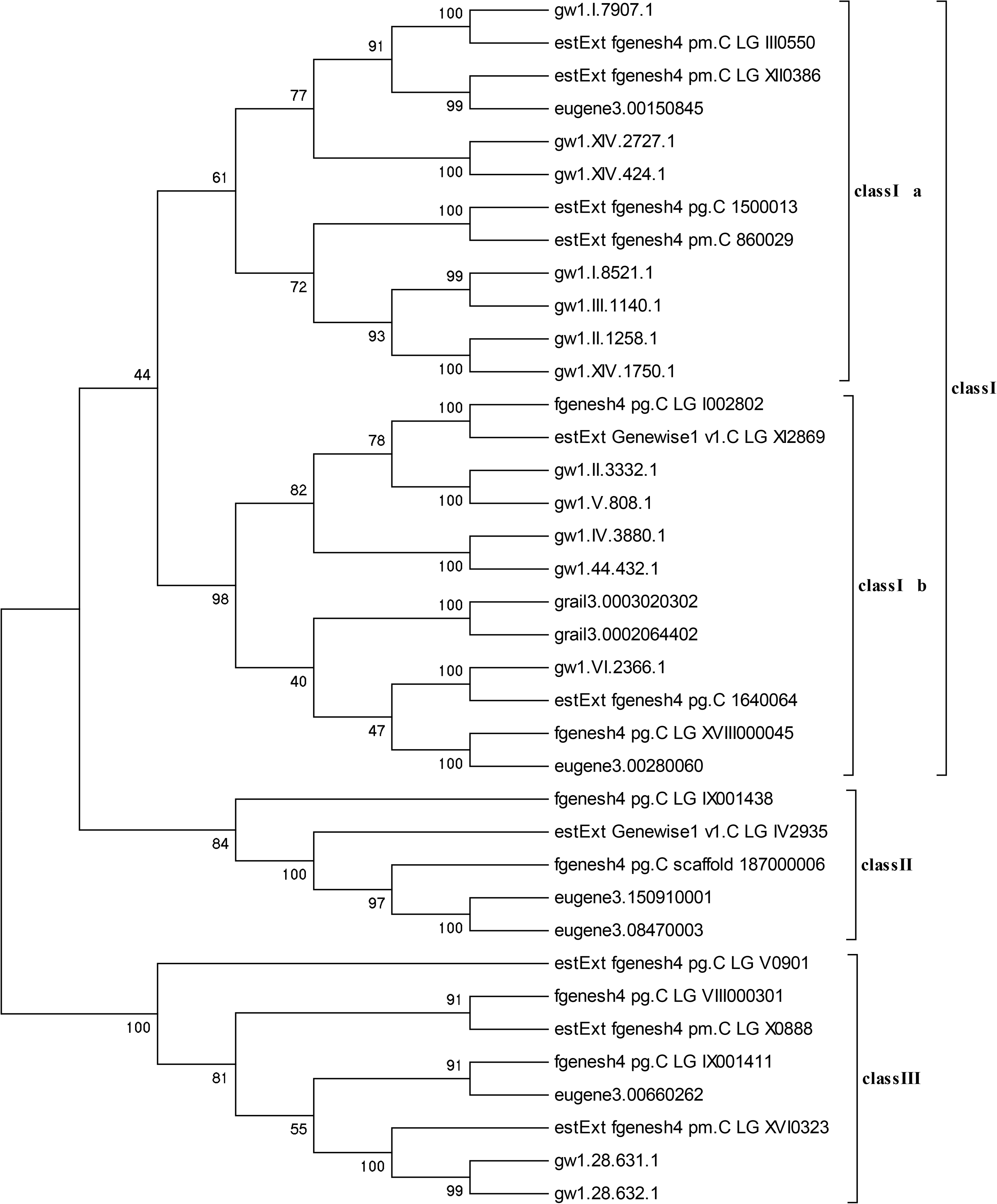
is the evolutionary tree made of MEGA4

Among the 37 ARF gene models, there is no corresponding probe representation for genes grail3.0003020302, grail3.0002064402, gw1.28.631.1, estext_fgenesh4_pg.c_1500013, and gw1.28.632.1, so these genes are not involved in our subsequent research and analysis.After Wilcox [4] [] test of PM and MM was conducted to determine whether the gene was expressed (gene expression was defined when p-value <0.05), only gene gw1.xiv.2727.1 was not expressed, and this gene was also excluded in the following study.

We conducted homologous and evolutionary analysis on the 31 ARF gene CDS to be studied, and the results are shown in figure [see figure 3.8]. After analysis, the 31 ARF genes can still be divided into three categories, the homology of classIa and classIb is increased to 54%, and the homology among the genes is generally improved, the lowest is 62%, making the homology and evolutionary analysis results credible.

**Figure 3.8.**
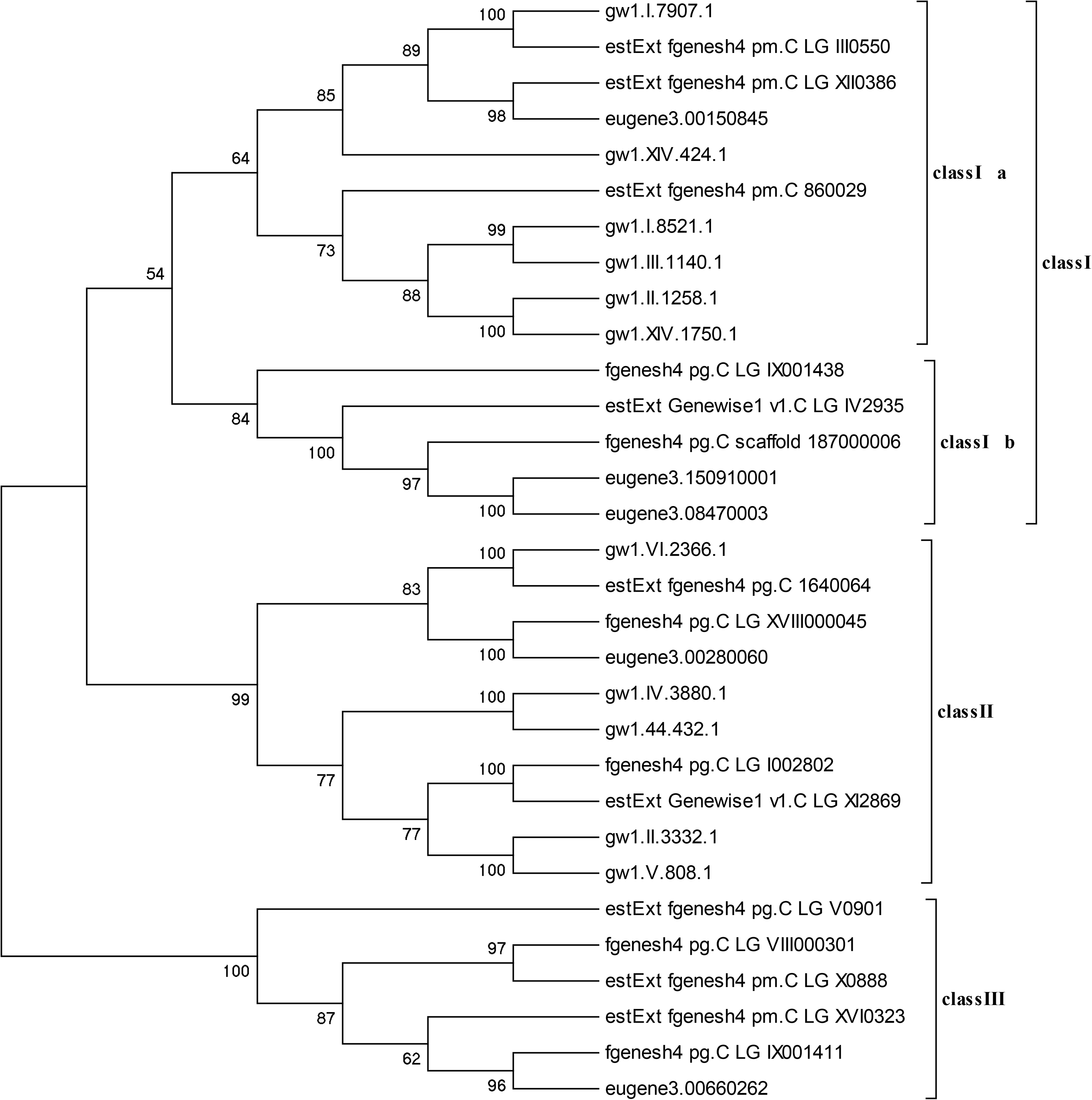
homologous and evolutionary analysis on the 31 ARF gene CDS

##### 3.4.2.2 correlation analysis of ARFs gene expression

According to the corresponding probes of ARF genes in the database, correlation analysis was conducted on the expression levels of the 31 ARF genes to be studied, and a network diagram of 31 ARF genes was made according to the correlation between different genes. When calculating the correlation coefficients of all expression levels, we chose the absolute value of correlation coefficients not less than 0.8 as correlation; otherwise, it was irrelevant, and drew the network diagram between 31 ARF genes [figure 3.9].

According to the analysis of 31 ARF gene homology evolution diagrams, gw1.i.8521.1 and gw1.iii.1140.1 genes have up to 99% homology, and the expression level correlation is greater than 0.8. These two genes have high homology and expression level correlation. The homology between eugene3.150910001 and eugene3.08470003 was 100%, and the intergene expression correlation was greater than 0.8.Estext_genewise1_v1.c_lg_iv2935, fgenesh4_pg.c_lg_ix001438 and fgenesh4_pg.c_scaffold_187000006 had the lowest homology of 84%, and the correlation coefficient of intergene expression was greater than 0.8.Estext_fgenesh4_pg.c_1640064, gw1.vi.2366.1 and eugen 3.00280060 had the lowest homology of 83%, and the correlation coefficient of intergene expression was greater than 0.8.Gw1.44.432.1 had 100% homology with gw1.iv.3880.1, and the intergene expression correlation coefficient was greater than 0.8.Gw1.V.808.1 had 100% homology with gw1.44.432.1, and the intergene expression correlation coefficient was greater than 0.8.Eugene3.00280060, fgenesh4_pg.c_lg_ix001411 and eugene3.00660262 all have high correlation coefficients with other genes, but the homology between these three genes and other genes is not high, that is, they are distributed in different species.Estext_fgenesh4_pm-c_lg_x0888 has a high correlation coefficient with the expression level of other genes, but the homology of estext_fgenesh4_pm-c_lg_x0888 is not high.

In conclusion, when intergene homology is higher than 80%, the correlation coefficient of intergene expression level is higher than 0.8, and intergene expression regulation may occur simultaneously. When the correlation coefficient of intergene expression level is higher than 0.8, the homology of genes is low and they are in different classes, which cannot predict the regulation mechanism of intergene expression. In other words, in future studies, we can determine the expression regulation mechanism among different genes according to the homology between genes and the correlation of gene expression.

#### 3.4.3 up-regulated and down-regulated genes of ARFs were selected

We used the median of 31 ARF genes corresponding to gene semaphores to determine up-regulated and down-regulated genes. When the gene expression increased by 80%, it was defined as up-regulated; when the gene expression decreased by 60%, it was defined as down-regulated; when it was in the middle, it was considered fine-tuning.

We determined the distribution of all up-regulated, down-regulated and fine-tuned genes in different tissues and seasons. The results are shown in figure [figure 3.10].

**Figure 3.10.**
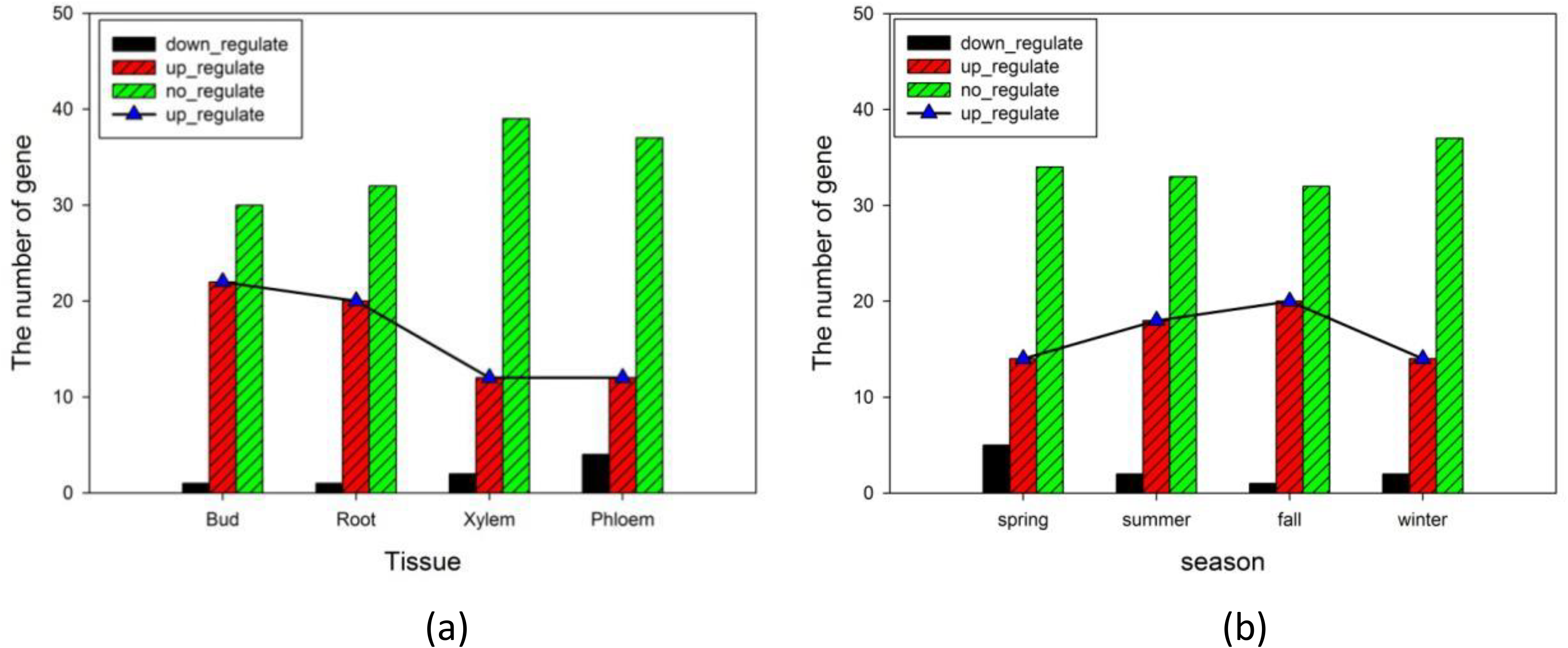
(a) up-regulated, down-regulated and fine-tuned gene profiles among different tissues; (b) up-regulation, down-regulation and fine-tuning of gene distribution in different seasons

Among the different tissues, the buds had the most upregulation genes, followed by the roots, and the xylem and phloem had the least upregulation genes. During plant growth and development, auxin is produced mainly by the terminal bud and transported to other parts of the plant through polarity. When auxin is produced, the auxin response factor can respond to auxin rapidly, and the number of up-regulated genes in buds is the largest. Meanwhile, the down-regulated genes are also less than other tissues. The number of upregulated genes was highest in autumn, followed by summer, with the same number in spring and winter.

We selected the distribution of up-regulated, down-regulated and fine-tuned genes in different tissues in the four seasons of spring, summer, autumn and winter.

In poplar bud tissue, the trend of upregulation of ARF gene semaphore in spring, summer, autumn and winter is shown in the figure 3.11. Semaphore gradually increases in spring, summer and autumn, and decreases in winter. The seasonal variation of auxin content measured in bud tissue is shown in the figure []. Both the solid line and the dotted line represent auxin content, but the coordinates are different. The content of auxin in bud increased first and then decreased along the four seasons of spring, summer, autumn and winter, and was the highest in autumn.

**Figure 3.11.**
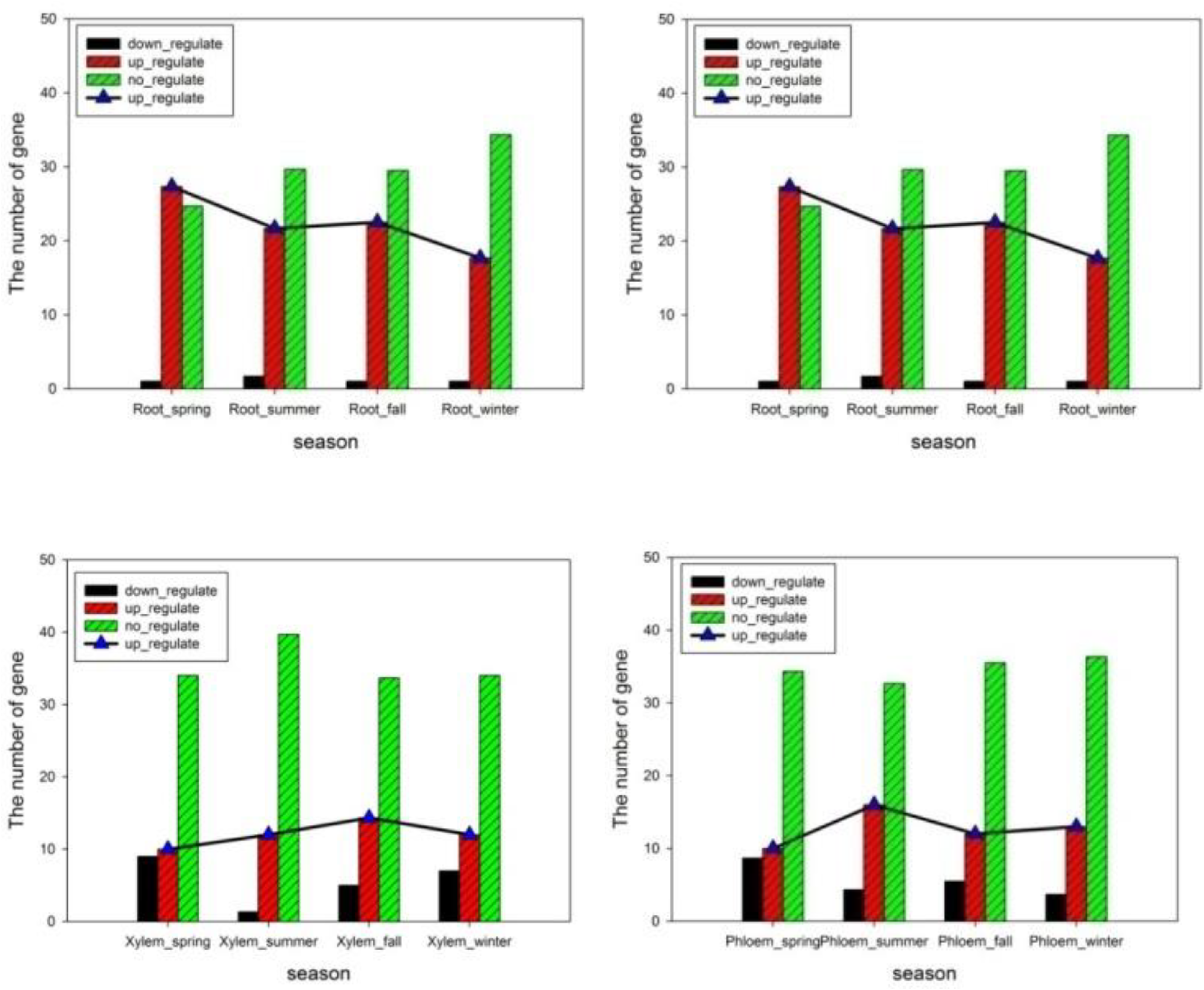
the trend of upregulation of ARF gene semaphore in spring, summer, autumn and winter

In Yang shugen tissues, the trend of up-regulated ARF gene semaphore in the four seasons is shown in the figure []. The root showed a downward trend with the up-regulated ARF gene in the four seasons of spring, summer, autumn and winter, with the highest in spring and the lowest in winter. The seasonal changes of auxin content measured in root tissues are shown in the figure []. It decreases in summer, increases in autumn, and decreases again in winter, among which autumn is the highest.

In the xylem tissues of poplar, the change trend of up-regulated ARF gene semaphore in the four seasons is shown. In xylem, the up-regulated ARF gene semaphore gradually increases with the spring, summer, autumn and three seasons, and slightly decreases in winter. The seasonal changes of auxin content measured in xylem are shown in the figure []. With the seasonal changes of spring, summer, autumn and winter, there is a declining trend, with the highest in spring and the lowest in winter.

In poplar phloem tissues, the trend of up-regulated ARF gene semaphore in spring, summer, autumn and winter is shown. In xylem, with the up-regulated ARF gene semaphore in spring, summer, autumn and winter, it first increases and then decreases, with the highest in summer. The seasonal changes of auxin content measured in phloem are shown. The solid and dotted lines all represent auxin content, but the coordinates are different. With the seasonal changes, auxin content increases first and then decreases, with the highest in summer and the lowest in winter.

We selected the up-regulated and down-regulated genes in different tissues and seasons, calculated the up-regulated or down-regulated multiples, and labeled the respective GO process and chromosome distribution [see table 3.6].

**table 3.6.** GO is only related to the growth pathway of auxin

| Gene | Chromosome | The GO | Up_regu<br>late | Down_reg<br>ulate |
| --- | --- | --- | --- | --- |
| Fgenes4_pg.<br>C_scaffold_187000006 | scaffold_187 | GO: 0009725 | --- | 0.616 |
| eugene3.150910001 | scaffold_15091 | GO: 0009725 | --- | 0.639 |
| eugene3.08470003 | Scaffold_ 847 | GO: 0009725 | --- | 0.639 |
| Gw1. II. 1258.1 | LG_II | GO: 0009725 | 6.12 | --- |
| Gw1. XIV. 1750.1 | LG_XIV | GO: 0009725 | 3.87 | --- |
| Gw1. VI. 2366.1 | LG_VI | GO: 0009725 | 12.8 | --- |
| EstExt_fgenes4_pg. C_1640064 | scaffold_164 | GO: 0009725 | 12.8 | --- |
| EstExt_fgenes4_pm. C_860029 | scaffold_86 | GO: 0009725 | 5.52 | --- |

It was found that estext_fgenesh4_pm.c_lg_xii0386 gene located on LG_XII chromosome showed both up-regulation and down-regulation in the same tissue in the same season.We were able to identify two different probe sequences for this gene using chip data provided by Affymetrix. After sequence homology comparison analysis, the homology of the two is less than 50%, we can infer that estext_fgenesh4_pm. C_LG_XII0386 contains different family members, whose homology is very low and expression level is different, either up-regulated or down-regulated.

#### 3.4.4 analysis of the relationship between the number of up-regulated ARFs genes and their semaphores and auxin content in each tissue

Auxin plays an important role in plant growth and development, and can promote plant growth cycle morphological changes, including cell division, differentiation and elongation, flower and vascular tissue development, as well as directional growth [40].The potential role of auxin as a morphogenetic promoter has been predicted in the formation of lateral organs. Plant growth hormone can promote the formation of secondary axes and the formation of tissue primordia from leaves to flowers [41–44].Auxin accumulation at these sites is accomplished by input and output transporters, whose activities make auxin in a dynamic flow state [43, 44].

Auxin promotes cell division in the middle column sheath, resulting in cambium [45–47].Auxin can regulate the changes of later plant organs, including the formation of pistil [48], leaf primordium, root tip [43] [] and other tissues and organs. In addition, in arabidopsis thaliana, Cheng [49] research prove that YUCCA flavin single oxidase gene in synthesis of auxin is required, it can adjust arabidopsis embryos and leaf form, and show that regulating auxin YUC family gene expression in the process of synthesis is the number of gene expression in the add and sex, the more the number of YUC family genes into plants, the higher of synthesis of auxin content. Cheng[49] et al. mainly conducted experiments with four genes in the YUCCA gene family, and the results showed that the cumulative expression of YUC gene could cause the excessive production of auxin.

In addition, Yoon[50] et al. concluded that the stimulation of exogenous auxin in lateral root tissue of arabidopsis thaliana could promote the up-regulation of ARF3 gene in lateral root, and within a certain range, the expression level of ARF3 increased with the increase of exogenous auxin concentration. In the experiment, exogenous abscisic acid was also used to treat the lateral roots, resulting in its inhibition of ARF3 expression. Even if the ARF3 gene expression was down-regulated, after the mixture of exogenous auxin and abscisic acid, the expression of ARF3 in the lateral roots was up-regulated compared with that in a single abscisic acid treatment. Therefore, it is inferred that the auxin response factor gene of poplar can accumulate and express, thus causing the excessive production of auxin.

We speculated that in poplar, more ARF up-regulated genes in different tissues could lead to higher auxin content in the tissues. To this end, we compared the changing trends of various organizations in the four seasons of spring, summer, autumn and winter(figure 3.12).

**Figure 3.12.**
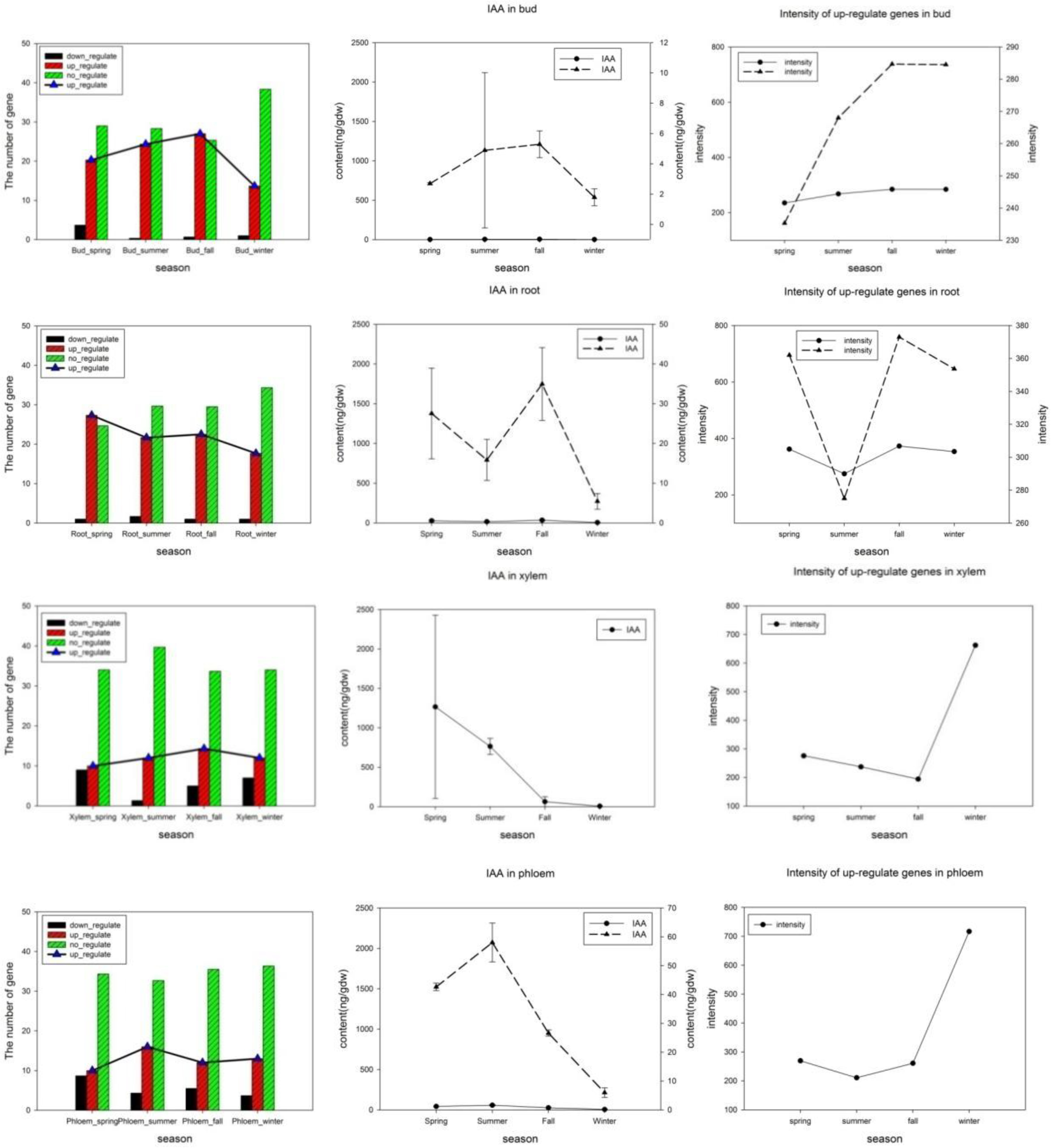
trends of various organizations in the four seasons of spring, summer, autumn and winter

In the four seasons of spring, summer, autumn and winter, the correlations among auxin content, gene expression semaphore and up-regulated gene number were calculated respectively in different tissues. It can be seen from the results that the correlation coefficients between gene expression semaphore and auxin content and up-regulated gene number are low, and there is no correlation between them. However, auxin content was highly correlated with the number of up-regulated genes in bud, root and xylem tissues, except phloem tissues. According to the correlation between auxin content and the number of up-regulated genes in four seasons, it can be concluded that in budding and root tissues, auxin content and auxin content show a high positive correlation, and a high negative correlation in xylem. The results showed that there was a correlation between the up-regulated gene data and auxin content in the same tissue, which was consistent with our predicted results.

It can be concluded that in any tissue(Table 3.7,Table 3.8,Table 3.9, Table 3.10), the correlation between auxin content, up-regulated gene number and gene expression semaphore pair is 1 or −1 in summer and fall, that is, highly correlated. During the process of poplar growth, the growth in summer and autumn is stable, which makes the stability of each research quantity and shows a high correlation.

**Table 3.7.** intensity/IAA/up-regulate, pairwise correlation analysis results of pairwise correlation in four seasons of different organizations between pairwise and pairwise, we defined correlation coefficient higher than 0.7 as correlation

|  | Bud | The Root | Xylem | Phloem |
| --- | --- | --- | --- | --- |
| Intensity/IAA | 0.236 | 0.417 | 0.441 | 0.862 |
| IAA/up - regulate | 0.959 | 0.718 | 0.801 | 0.316 |
| Intensity/up to regulate | 0.004 | 0.179 | 0.183 | 0.041 |

**Table 3.8.** correlation coefficient between gene expression semaphore and auxin content: (The correlation between intensity of up-regulation gene and The content of IAA)

|  | Summer/fall | Spring/summer/fall | Summer/fall/winter | all |
| --- | --- | --- | --- | --- |
| Bud | 1 | 0.981 | 0.399 | 0.236 |
| The Root | 1 | 0.958 | 0.348 | 0.417 |
| Xylem | 1 | 0.998 | 0.486 | 0.441 |
| Phloem | - 1 | 0.782 | 0.851 | 0.862 |

**Table 3.9.** correlation coefficient between auxin content and up-regulated gene number: (The correlation between The number of up-regulate gene and The content of IAA)

|  | Summer/fall | Spring/summer/fall | Summer/fall/winter | all |
| --- | --- | --- | --- | --- |
| Bud | 1 | 0.965 | 0.996 | 0.959 |
| The Root | 1 | 0.262 | 0.863 | 0.718 |
| Xylem | - 1 | 0.998 | 0.441 | 0.801 |
| Phloem | 1 | 0.646 | 0.798 | 0.316 |

**Table 3.10.**
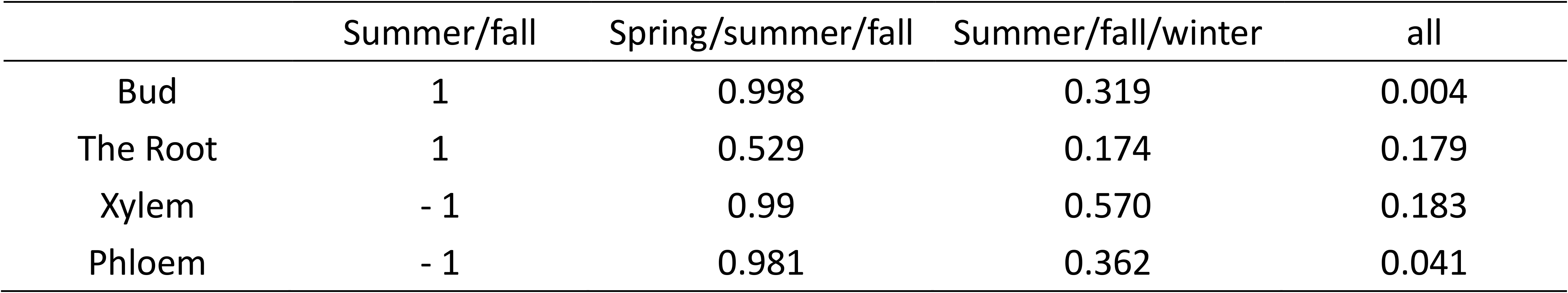
correlation coefficient between gene expression semaphore and up-regulated gene number: (The correlation between The intensity of up-regulate gene and The number of up-regulate gene)

However, once winter data were added, the correlation between gene expression signaling and auxin content and the correlation between gene expression signaling and the number of up-regulated genes decreased, but the correlation between auxin content and the number of up-regulated genes was not obvious, indicating that winter had a greater impact on the up-regulated gene expression signaling. Plant growth and development are affected by external environmental factors, including biological and abiotic stress and endogenous growth factors (i.e., plant hormones) regulation. In general, plants respond to the effects of the environment by changing the amount of endogenous hormones. Among a large number of abiotic stress, low temperature has a greater influence, which can limit the growth and development of plants [51] [].Under low temperature stress, abscisic acid plays an important regulatory role in promoting the rapid adaptation of plants to the maladaptive environment [52, 53]. Yoon[50]et al. proved that abscisic acid can inhibit the expression of medium ARF gene in plant root tips, which is contrary to the effect of auxin.Shibasaki[51] et al. treated the root tip of arabidopsis thaliana at 4°C. Through the influence of root tip gravity and root tip length, it was proved that low temperature could affect the transport of auxin to a greater extent, and had less impact on the transmission of auxin signal. Under low temperature stress, the content of abscisic acid and ethylene in plants increases, and the content of plant hormones such as jasmonic acid and salicylic acid also increases correspondingly [54]. In winter, due to the mutual regulation of various hormones and the restriction of auxin signal transmission, the results obtained from winter data were deviated greatly.

In the four tissues we studied, shoot, root, xylem and phloem, auxin is produced by shoot and transported to root and xylem through phloem according to the source bank theory of plants, promoting the growth and development of plants. In this process, plants also produce gibberellin, abscisic acid, cytokinin and other plant hormones, which regulate each other and jointly control the growth of plants, thus showing positive or negative correlations in different tissues.

## 4 look

Using the microarray data of poplar gene in the laboratory, the phloem and xylem genes with good clustering effect were selected for research and analysis after clustering evaluation. Different genes (p-value <0.05) were selected based on Wilcox non-parametric test, and a gene related to plant hormones and expressed was selected for analysis. After the prediction by NCBI, softberry, Pfam and other databases, the function of this gene was determined as auxin response factor.

Auxin response factor genes were selected from the transcription factor database of poplar, and evolutionary tree and relationship analysis were conducted on the ARFs family of poplar. For genes with less than 100% homology, the correlation coefficient of gene expression may be higher than 0.8 or lower than 0.8.

The median was used to select the up-regulated and down-regulated gene distribution in different tissues of poplar ARFs family in the four seasons, and then we made the distribution map of gene expression level in different tissues in the four seasons. The results showed that the variation of up-regulated gene number in different tissues with the change of the four seasons was the same as that of gene expression level.

The interaction between IAA and cytokinin, abscisic acid, gibberellin, ethylene and other hormones resulted in a great difference in the seasonal variation trend of IAA in the root. At the same time, we can use the transgenic RNA interference test for subsequent experimental verification of our results.

## Appendix

> consensus: Poplar: PtpAffx. 205505.1 S1_at; Pmrna | pmrna10882;

gtggagagggcaaagtgaaggtagaggagggattttcgatgagtgggagagggaggttgt

cccaggaggctgtggcggaggctgtggagatggcggctaaaggtttgccgtttgacgttg

tgtattacccgagggcaggttggtactcagattttgtggtgagggctgaagcggtggagg

ctgcccttggtgtgttttggactgctgggatgagagtgaagatggccatggagactgagg

attcttctagaatgacttggtttcaggggactgtttcgggtactggcttaccagactctg

gtgcatggcgtggctctccttggcgcatgcttcagattacgtgggatgagcctgaagttt

tgcagaatgcaaagagagtgagcccttggcaagttgaatttgttgcaaccactccacagc

ttcaagctgcattccccccaatgaagaaattgagatatccaaatgattctaggttcctga

cagatggagaacttttttttcccatgtcagatctaactaattcaactatggggcacacaa

atgcttcaatgttgaattatagtacttttcctgctggcatgcagggagccaggcaagatc

ctttttccacattcggtttatccaactttataagtgaaaatgctcctcaggtgttcagtg

acagggcctttggaaacaacttggtccctaagatgaaaagaatgcccagtgagatgaaca

ttggcagtttgcagtctgagaacttatcacctgagagccagagtagtgcatattcttttg

gcataggttttgttggaaacaggggcttcaaccccaagaaagttggaattaactctattc

agttatttggtaagatcattcacatggaccagcctgttgaaaatggttttgatgatgttg

gcttcatggatgacagcagtaaatgttgcaatgaaactgaaggcgttgatgcactggaac

tttcactgacttcctcctacacagaactacttaacaggattgatgcccaatgccaaagag

cttcacctgttgaggcatgtcctacttga

1|:10713019-10715735 Populus trichocarpa linkage group V, whole genome shotgun sequence

GAAAGCTGTATCATGTAGTTATAGTTGCTTCTTAATAGTAAAATACTTGCAAAACTTGTTATATATTAAA

CAGGAATTGTATTATCTGATTCCCAATTACATGATAAAATCAGGAGATTGGAAGAAATCCTTCGGAATGT

ACATCACCGAACTTTTGAATTTGAACAGATTCTTCTAGTATTGACCAGTAACCATATAGCATGATGCTTT

TACTACACAGCAATTCATAGAAATCCTTTGGAACTTTGTTCTGCTTCAAAACAAATAAATGGTTTTCACT

ATTTATTTCAGCACAAACTATATTAGACAAAGAATTAGTTCTTCAGAGGCATTTGACTGGCTGTAGATAT

CTTTGAAAATCAAGACCCAAGAAAAAAAGTAGTGTATAGTGTACCATGTTATGCTTCTGAAAGCCAAAGA

GCTGTTTATTTCAACTAATTTGATTGTCACTAGTATATTAGAACAGGACCACTTACTCCTCAATGGACAC

CATATTTACCTTCTAATGAAAAAAGAAACCTGCAGTTTATTTCAACAACAAACAGGTTTGCTTAGGAATA

AGTTAAAGAATGCAAAGATCTAAAGATATCTATAAATCAACGTAGATTCTGAACAAATGTAAGGAGAAGT

GTCAGAGGAATTGCTCACACTGGAACCACACTCTCTAAATCTAGTGCAAGATTAACATGTGCATGATACA

TTTAAACAAATTCCTAGGGAAGCATGATACATGAAAAGAGACATGACAACAATGTAATGACATTAAAGCT

AAATGAAGAACATTTGAGATAAGGCTCAATTAGTTGAAATTGCATGGCAGTAACCATATTATAGGAACTT

GATCATTAATTAAGTATTTGTAAACAACTAACTAAGATGCACAATCAAATTTAAAGAAGTAACATTTCAG

GCAAGTTTAAGATAATAGAGTTTGTGGGGTCCAATCACAAAAAACATACCTGCACAGCTACTTATATGCA

TGCGCTACAGAACTACTTGTTCAAGTAGGACATGCCTCAACAGGTGAAGCTCTTTGGCATTGGGCATCAA

TCCTGTTAAGTAGTTCTGTGTAGGAGGAAGTCAGTGAAAGTTCCAGTGCATCAACGCCTTCAGTTTCATT

GCAACATTTACTGCTGTCATCCATGAAGCCAACATCATCAAAACCATTTTCAACAGGCTGGTCCATGTGA

ATGATCTTACCAAATAACTGAATAGAGTTAATTCCAACTTTCTTGGGGTTGAAGCCCCTGTTTCCAACAA

AACCTATGCCAAAAGAATATGCACTACTCTGGCTCTCAGGTGATAAGTTCTCAGACTGCAAACTGCCAAT

GTTCATCTCACTGGGCATTCTTTTCATCTTAGGGACCAAGTTGTTTCCAAAGGCCCTGTCACTGAACACC

TGAGGAGCATTTTCACTTATAAAGTTGGATAAACCGAATGTGGAAAAAGGATCTTGCCTGGCTCCCTGCA

TGCCAGCAGGAAAAGTACTATAATTCAACATTGAAGCATTTGTGTGCCCCATAGTTGAATTAGTTAGATC

TGACATGGGAAAAAAAAGTTCTCCATCTGTCAGGAACCTAGAATCATTTGGATATCTCAATTTCTTCATT

GGGGGGAATGCAGCTTGAAGCTGTGGAGTGGTTGCAACAAATTCAACTTGCCAAGGGCTCACTCTCTTTG

CATTCTGCAAAACTTCAGGCTCATCCCACGTAATCTGTGCAACAGAGAAAAAACAAGTTGAAAAGGCAGA

AAACACAGTAAAAAGATCAAAAGTTTTCACTCTGTTATAATGAAGAAATTTCTGAATGTTAAAGCTTATA

TATCAGCAACTATATAACTGAATCATTAACAGATAGTCACTTGTGAAGCTCATTACTTGACCTAAAACTT

TTAGACACTTGAGAACTAAGAAGATTGAAAGACCTGGTTAAAATTACTTAACCTTCACAATCAAACTGCA

GACATTGTGAAAACCTAGACATAAATAAAATCAGAAAGAGGAACAGCATGAAAAAGATGTAAGAGAGTTA

AAACAGCATACGCTGATATCCAATTTGCATGTGCAATTGAATCTAGTTCTAACAATATTCTTAATTGAAC

CCCAAACTATTATAACATCTGCCTGACCTAGCTTTCTTGACATTGGGCAAATAAAATGGCCCCTATGAAA

GGGCAAATGGCTCATGGAACAAATCAGAAGTCCTTAACAAATTCCATAGCGATATTCTAATTTCAATGAA

AACCAAGTTCTTAAAACTTTAAGTGGTACTCCTCCACAAGGAATTGAAGAAAACCTCAATCACGTTTCAT

AACATATAGCTTGCAGATAAACAACGTAGCCTGGGGTAGAATGAAGAACAAAGACAAAAAAGGTACGACA

CAGTAGCTTAGCTGCACTACACTGCCCTGAAAACTCAATACCCAAAGCACCAGAGCCACAGGAAACAGCA

CACAAATAAAATTAAAAAAACGGGGAAACTGTGAAAAGAATACCACTATTTCAAGCATCACCCAGAATCA

GATATCCAATTTATAAAATACATAGCTCCATTAGCACCAACTGTTCTTCATTTTCTCTGGCATCGTTTTC

ATTGATTCTTAATGATTAATATACACTCCAGATATCATTAGCAAAAAATCGCAACAGGGGAGATTATCCC

AAAGCATTTTTCTGTTCAAAGTCAAAATTAGCAGGTAATGTTTAGTGCATATGCAAG

GPDNN algorithm model.

1. **Wilcoxon symbol test was used to determine specific binding and non-specific binding based on p value** Training data set. Arrange the p-values from small to large, and take the first 1000 probes with a p-value less than 0.05 Set was the expression probe set, and the PM probe sequence and intensity of the gene were trained as specific binding parameters Set. Randomly select 1000 probes from a set with a p value greater than 1/4 or median The PM and MM probe sequences and intensities are nonspecific binding Training set of parameters.
2. **On the training set without expression of probe set, its specific binding strength was zero, and there was no specific binding** Target sequence. Formula to

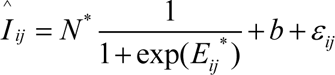 By minimizing the function

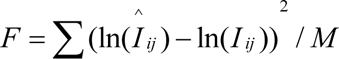 The most rapid descent method was used to determine the non-specific parameters, including position parameters w and free energy parameters e (bk);*Bk +1) and N¤and b*.

